# *micromorph*: a Python toolkit for measurement of microbial morphology

**DOI:** 10.64898/2025.12.05.692496

**Authors:** Simone Coppola, Séamus Holden

**Affiliations:** School of Life Sciences, University of Warwick, Gibbet Hill Campus, Coventry, UK

## Abstract

Detection of morphological phenotypes from light microscopy is a key part of microbiology. Despite advances in automated morphological analysis, accurate measurements still often require significant user input. To address this, we have developed *micromorph*, a Python package to measure bacterial morphological properties from multiple types of light microscopy data. *micromorph* is available as a Python package with a documented API, or as *napari* plugin, thus appealing to a range of users regardless of their coding proficiency. The *micromorph* API allows easy integration into custom analysis pipelines. We demonstrate a range of scenarios in which *micromorph* demonstrates excellent performance and introduce a new approach for reliably measuring the widths and lengths of quasi-circular cells.

## Introduction

Morphological phenotypes, such as changes in cell shape or size, provide important insights into gene function and bacterial responses to external stimuli (1–3). These changes are often subtle or may be hidden by variations in shape within each sample, requiring reliable and accurate measurement methods to detect them. Furthermore, advances in high-throughput microscopy (4) have significantly increased data throughput in recent years, making automated analysis methods essential to enable fast, reliable and unbiased measurements.

Several tools have been developed to automate bacterial morphology analysis (5–8). Although these packages have proven to be powerful tools for bacterial cell biology community, they have drawbacks. *MicrobeJ* and *Oufti* are all-in-one solutions capable of analysing bacteria with a wide range of morphological types, but they have been optimised for phase-contrast microscopy images, making them unsuitable for chaining cells. *ChainTracer* is better suited for chaining bacteria such as *B. subtilis*, but it is not fully automated, requires both phase-contrast and membrane-stained images to correctly measure widths and does not perform well with non-chaining cells. Furthermore, all the above mentioned software expose only a limited Application Programming Interface (API), making them complex to be integrated into custom analysis pipelines. This also makes them challenging to integrate with the latest Python-based deep-learning segmentation algorithms, which have been shown to outperform classical image processing approaches in segmentation algorithms (9–11). Surprisingly, although Python is the most widely used language for machine learning, there is currently no software capable of precise morphology measurements of complex cells shapes that runs natively in Python, with packages such as *RabitPy* and *TrackRefiner* instead opting to use the length of axes of a fitted ellipse to the mask as a proxy for width and length (12,13)

Here we present *micromorph*, a Python package for automated analysis of bacterial morphology designed to address the limitations described above. *micromorph* can automatically quantify morphology for phase-contrast images and for fluorescence images of cell envelope or cytoplasm stained bacteria. Specifically: (i) deep-learning-based image segmentation is provided via Omnipose (10) or users can import masks from other software; (ii) three morphology analysis algorithms are implemented, capable of robust and precise measurement of cell dimensions for diverse cell morphologies and multiple microscopy modalities; (iii) a graphical interface to the software is provided via a *napari* plugin (14)

A full API with supporting documentation is provided with the aim of making *micromorph* suitable for users aiming to either customise parts of the analysis or integrate the analysis in custom pipelines, such as, for example, using a different segmentation algorithm instead of Omnipose.

In this paper, we demonstrate analysis of a variety of datasets with *micromorph*, with a particular focus on highlighting the flexibility of the software for different bacterial morphologies and imaging modalities. We demonstrate improved performance compared with other software packages. We also introduce a novel method for measuring cell widths and lengths in coccoid or otherwise rounded bacteria.

## Methods

### Cell culture and staining

*B. subtilis* strains were streaked from -80°C freezer glycerol stocks onto Nutrient Agar (NA) plates containing the appropriate antibiotics and grown overnight at 30°C. Starter cultures were inoculated from a single colony and grown overnight on an orbital shaking incubator (175rpm, 30C) in 2ml LB + 2% glucose. The following day, cultures were diluted to OD600 0.1 in 5ml LB and grown until OD600 reached 0.5-0.7.

For membrane staining with Nile Red, at that point, cells were diluted back to OD600 0.1 in 200ul LB containing Nile Red (1ug/ml final concentration) and incubated for 10 minutes at 30°C in a shaking heat block (ThermoFisher). For cell wall staining with sBADA staining, cells were instead diluted back to OD600 0.1 in 200ul LB with sBADA (50uM final concentration) and incubated for 30 minutes in a shaking heat block.

Cells were washed in fresh, pre-warmed LB twice before being concentrated 50x for final use.

The same protocol was followed for the preparation of wild-type *E. coli* (MG1655) and *Staph. aureus* (ATCC 29213) cells. In those cases, all incubations took place at 37°C instead of 30°C.

For the *ftsW* knockdown experiment, overnight cultures of BEC14850 (*trpC2, lacA::Pxyl-dcas9(ErmR), amyE::Pveg-sgRNA_ftsW(CmR))* were grown in LB + 2% glucose as described above. In the morning, cultures were diluted 1:200 and spotted on a flat Nutrient Agar pad with added 0.1% and left to grow at room temperature for 24 hours after enclosing the pad with the coverslip (see below), after which the phase-contrast images were acquired.

### Agarose pad preparation

Flat 2% agarose pads in 1:10 LB were prepared by pipetting 50ul of molten solution inside Geneframes (Thermo Scientific). A 1ul droplet of the final cell suspension was then spotted on the flat pads. The droplet was then left to dry for one minute before placing a 22×22mm coverslip on top of the pad for imaging (15).

### Microscopy

Microscopy images used for initial testing (Figure 2) were acquired on a Nikon Eclipse Ti-2 microscope equipped with a 100x Ph3 objective (Nikon CFI Plan Fluor DLL 100X Oil). An Andor Zyla sCMOS camera was used to acquire images at a final pixel size of 65 nm/px. Phase contrast imaging was achieved by illuminating the sample using a 525nm LED (pE-100, coolLED) through a condenser equipped with a Ph3 phase annulus. A multi-colour LED engine provided illumination for fluorescence microscopy. For each position, images were acquired sequentially (phase, GFP, FM4-64), with a quad-band dichroic mirror and filter set (DA/FI/TR/Cy5-A-ZHE, Semrock) in the optical path.

### Synthetic data generation and analysis

Synthetic microscopy images (Supplementary Figure 3) were generated using the *microsim* package for Python (16). In short, a ground truth mask with the desired morphological properties is fed through an algorithm which convolves it with a point-spread-function matching that of a specified fluorophore being imaged through a microscopy system. Noise was added to the images using the *random_noise* function from the *scikit-image* package, with a noise sigma of 0.001 (17). All code used to generate the images is available in the *micromorph* Github repository, and samples from the synthetic image datasets have been included in the Github repository and Zenodo release (https://doi.org/10.5281/zenodo.19386228). Masks for image analysis where generated using *Omnipose* after training the *bact_fluor_omni* model further on a set of 500 synthetic images (which were not used for the benchmarks afterwards). We have included the model in the Zenodo release. We did not generate a phase-contrast synthetic dataset since the cytoplasmic fit is the inverse of the phase-contrast one.

### Implementation and data analysis

All code and installation instructions are available on GitHub (https://github.com/HoldenLab/micromorph/). An environment matching the one used for testing can be generated using Conda, or the package can be installed directly in *napari* via the plugin manager. All computations were performed on a Dell workstation equipped with an Intel i9-13900 processor and 64GB of memory. For the *MicrobeJ* analysis (5), data generated in Fiji (18) was first exported in the .mat format and converted to .csv using custom MATLAB scripts (Mathworks, R2023b). Figure panels were generated using custom Python scripts and assembled into complete figures in Inkscape (https://inkscape.org/).

## Results and discussion

### micromorph is a Python package suitable for a wide range of users

*micromorph* can measure bacterial cell boundaries, medial axes and other morphological metrics, with a specific focus on the accurate measurement of cell widths and lengths for a wide range of cell morphologies, including rod-shaped, spherical and elongated. *micromorph* uses the *Omnipose* deep learning based segmentation software (10) to obtain cell outlines from microscopy data for downstream morphology analysis. *Omnipose* comes with two built-in models for bacterial segmentation which we have found to perform well on our datasets. Alternatively, users can tailor the *Omnipose* models for their data by transfer learning, or import masks generated by other segmentation tools for analysis with *micromorph* alongside the raw microscopy data.

*micromorph* can be utilised in multiple ways, depending on user expertise and aims (Figure 1). The *napari* plugin is suitable for users requiring a simple installation process and a GUI based experience. As well as being useful for optimising parameters and visualising results, the *napari* plugin features a batch utility to automate the analysis of multiple datasets. Users interested in integrating *micromorph* into custom analysis pipelines can instead use the provided API to access relevant functions. All code and documentation are available on GitHub. There, we also provide installation details and example scripts.

**Figure 1.**
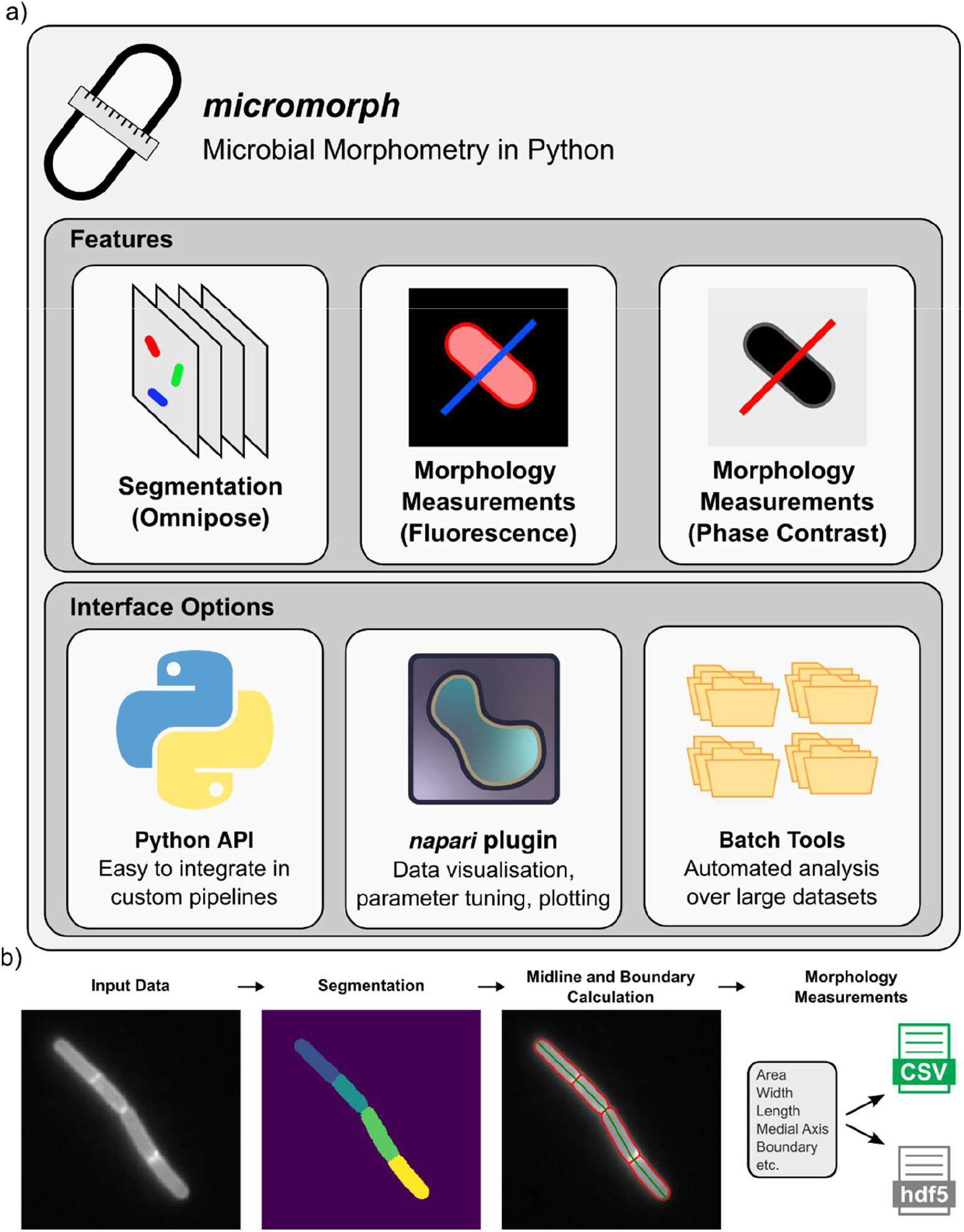
a) Overview of micromorph features. micromorph allows users to obtain morphological measurements from either phase contrast or fluorescence microscopy data. Functionalities can be accessed through a Python API or a napari plugin. A batch tools interface is also provided to automate analysis of multiple folders. b) example workflow and outputs for micromorph.

### micromorph can find widths and lengths of bacteria imaged through different modalities

Many popular bacterial morphology analysis tools measure cell widths by extracting this parameter from the fitted boundary of the bacteria (5,19). While this approach can work well, it has certain limitations – such as relying on good segmentation and being dependent on user-defined smoothing parameters.

A different approach involves measuring the intensity profile of the image perpendicular to the long axis of the bacteria at various (or one) points along its length and fitting it with appropriate functions depending on the imaging modality with which the image was acquired. This approach is particularly useful when segmentation results are not optimal, as it only uses the initial boundary approximation to calculate the direction of the lines along which the intensity is measured. While popular, previously this approach has only been implemented in a semi-automated fashion (1,2,4). In these cases ROIs from which to extract intensity profiles have been manually drawn to overcome limitations posed by threshold-based segmentation methods. This naturally introduces constraints on throughput as well as potential bias.

In *micromorph*, we implemented automated intensity-based estimation of cell width from multiple line profiles along the cell’s medial axis. Furthermore, we developed three different fitting models to cover most imaging scenarios typically encountered in bacterial imaging: phase contrast microscopy, fluorescence microscopy of membrane stained bacteria and fluorescence microscopy of bacteria expressing cytoplasmic fluorescent proteins (Figure 2a, b, c). Details of the fitting methods are described in the Supplementary Materials.

**Figure 2:**
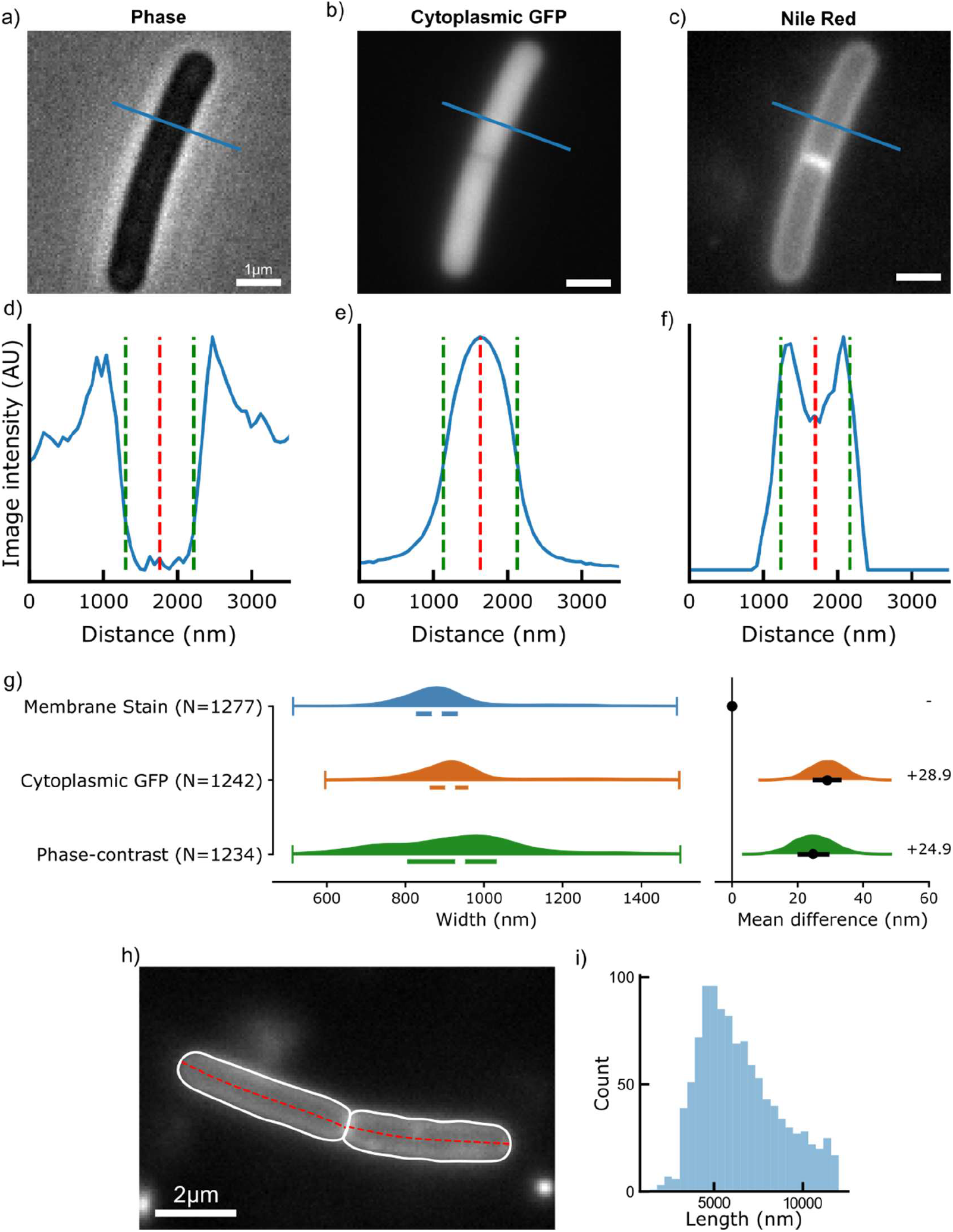
a, d) representative phase contrast image and fitted profile, green dotted line represents cell edges, red dotted line is the middle of the cell. b, e) same for cytoplasmic GFP cell and (c, f) membrane (Nile Red) stained cell. g) Left: comparison of widths extracted from ∼1300 cells. Median values are: 917nm, 880nm, 944nm for membrane stained, cytoplasmic GFP and phase contrast images respectively. Right: DABEST (20) plots showing effect size analysis. h) cell boundary and midline as calculated by micromorph. The midline is used to determine cell length. (i) cell length histogram from the membrane-stained dataset.

To validate our width finding algorithms, we acquired microscopy images in all three imaging modalities for a cytoplasmic GFP-expressing strain of the monoderm *B. subtilis* which was also membrane stained with Nile Red (see *Methods, Cell Culture and Staining*). Despite the large differences in the cell line profiles for different image modalities, we found that the three methods yielded consistent width measurements with only small shifts between the means of each method (Figure 2g). To quantify this, we analysed the data using estimation statistics (20), showing that there is a <30 nm difference in cell width measurements, regardless of fitting method (28.9nm [17.9, 39.4 95%CI] for membrane-stained minus cytoplasmic and 24.9nm [13.4, 36.0 95%CI] for membrane-stained minus phase-contrast). Further to this, analysis from 1000 synthetically generated cells showed our membrane fitting algorithm to have a mean difference of 65.8nm [64.4, 67.0 95%CI] from the ground truth, while our cytoplasmic/phase-contrast fitting algorithm had a difference of -67.8nm [-69.8, -65.2 95%CI], as well as being robust to fitting in the case of low signal to noise ratio images (Supplementary Table 2, Supplementary Figures 4 and 5).

We also implemented length measurement in *micromorph*. We make use of the interpolated cell boundary to calculate the cell medial axis and then extend this medial axis along the whole cell length. We follow this with a series of interpolation steps until the error between two consecutive interpolations is minimized (Figure 2h). The length is then measured to be the total arclength of the cell midline, demonstrated here by measuring the length of membrane-stained cells described above (Figure 2i). Similarly to the width finding algorithm, our length finding algorithm was found to be accurate with a mean difference of 90.2nm [88.8, 91.6 95% CI] from the ground truth data (Supplementary Table 2, Supplementary Figure 4).

### Line profile morphology estimation plus Omnipose segmentation enables accurate analysis of high-density bacterial images

Cell density is a frequent obstacle in image analysis. Thresholding algorithms can generally successfully segment bacteria from background, but usually need further ad-hoc processing to correctly distinguish adjacent cells. This makes the analysis of microcolonies or clumps of cells complex, unreliable or needing user adjustment. Deep Learning-based algorithms have been able to overcome this obstacle (11,21), opening up the possibility to more reliably analyse high cell density images.

High cell density can cause problems even in the case of semi-manual analysis. In the case of a membrane-stained cell, if a line profile drawn across a cell goes through part of a second cell, the intensity profile may show three peaks instead of two, or a skewed profile. Similarly, the intensity profile might not show a clear top-hat profile to fit in the case of phase contrast images. This leads to unexpected and/or incorrect results, particularly for automated analysis pipelines.

Accurate segmentation allows us to overcome this limitation. Specifically, we apply the mask to the original image before measuring the intensity profile, i.e. we replace any pixel value outside the cell with the calculated image background value. We then apply our width/length fitting routines to this adjusted intensity profile. Figure 3(a-f) illustrates the difference between applying or not apply the mask in the context of analysing phase contrast images of wild-type *E. coli* (MG1655). When the original image as the basis for the intensity profile, the fit fails and the measured width is 1899 nm (Figure 3c). When the processed image is used as a basis for the analysis, the width is measured to be 1040 nm (Figure 3f). By including information from *Omnipose*’s accurate segmentation in the analysis, *micromorph* can measure cell morphology in both low and high cell density images. We demonstrate *micromorph*’s performance in high density images by measuring the width and length of *E. coli* (MG1655) cells in a microcolony, using a test dataset available from (22) (Figure 3g-i). This method also improves width finding of membrane-stained cells (Supplementary Figure 1). We confirmed the usefulness of this approach by testing it on synthetic data, showing that mask application prior to fitting is critical for accurate measurement of dense cytoplasmic labelled cells, as well as yielding improved results for membrane labelled cells in close proximity (see Supplementary Table 2, Supplementary Figure 7).

**Figure 3:**
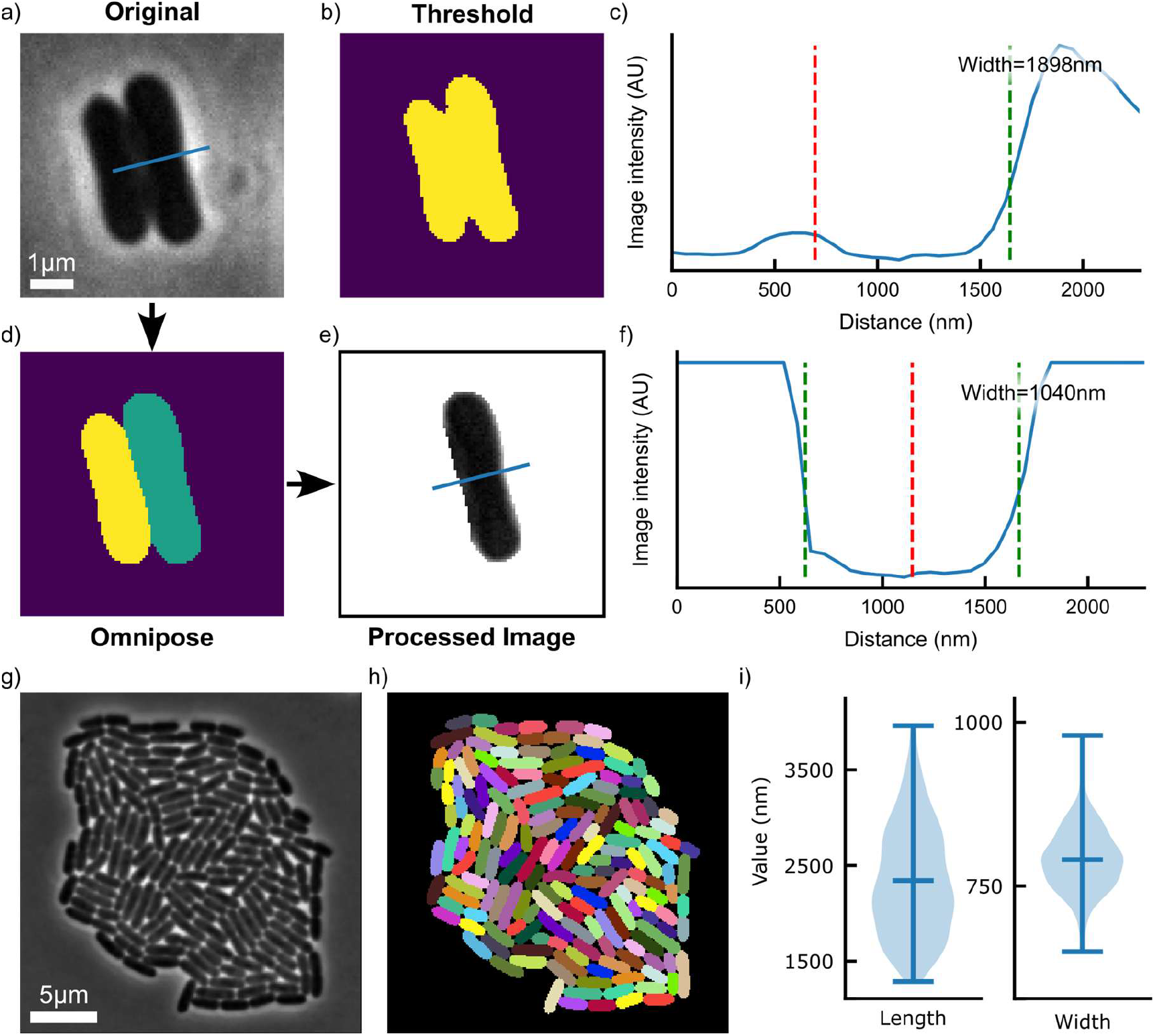
(a-f) Example where integration of segmentation information and intensity analysis enables robust morphology analysis. Two E. coli cells (MG1655) are clumped together. Top panels: the intensity profile is extracted from the original image, and the measured width is incorrect. Bottom panels: the intensity profile is extracted from the image after mask application, and the width is calculated correctly. Red dotted lines indicate centre of fitted profile, green dotted lines indicate cell edges from fitted profile. The second line from the top panel is not visible due being well out of the x-axis bounds. (g) Example phase-contrast image of an E. coli (MG1655) microcolony from (22) and the corresponding segmentation from Omnipose (h). (i) Width and length analysis for the cells in the microcolony.

### Measure360 enables reliable measurements of non-rod-shaped bacteria

When measuring the width or length of a bacterium, the first task is typically determining its short and long axes. In default mode, *micromorph* measures widths by first finding the midline of the cell starting from a medial axis transform (23) which is then optimised and smoothed. Widths are then calculated by fitting intensity profiles perpendicular to the midline. It therefore follows that accurate width measurements are only possible with this method if the midline is correctly defined at the start of the process. This approach, henceforth termed the “ShortAxis” algorithm, is well suited to dealing with rod-shaped cells, where a correct midline can be reliably defined starting from the medial axis transform.

The ShortAxis width measurement algorithm becomes more complex if cells have an approximately circular or elliptical shape. In such cases it might be impossible to define a midline (the medial axis transform of a circle is a single point) or the midline may not be accurate, thus leading to higher values of the width being measured (Figure 4a).

To overcome this issue, we developed the “Measure360” algorithm: an alternative way to obtain intensity profiles to be fitted to measure widths and lengths. In this case, the only information required for measuring widths is the centroid of the object of interest, which can be automatically obtained from the segmented cell mask. Intensity profiles are measured over multiple angles covering the 360 degrees around the centroid (Figure 4A). By measuring the interpeak distance at each angle it is possible to determine an interpeak distance profile around the entire object. The data are then smoothed to remove any outliers. After that, the width and length of the object are defined as the smallest and largest distances in the profile respectively (Figure 4c). Measure360 is suitable for either rod shaped or circular cells, with its main limitation not being suitable for highly curved cells, where the ShortAxis approach is recommended. We confirmed this by analysing three sets of synthetic data, where cells had either a half-moon, rod or elongated shape (see Supplementary Figures 7 and 8). Our analysis showed the two methods had comparable performance on rod-shaped cells, but Measure360 performed better on the half-moon dataset while the ShortAxis method performed better on the elongated cells (see Supplementary Table 2 for full details).

**Figure 4:**
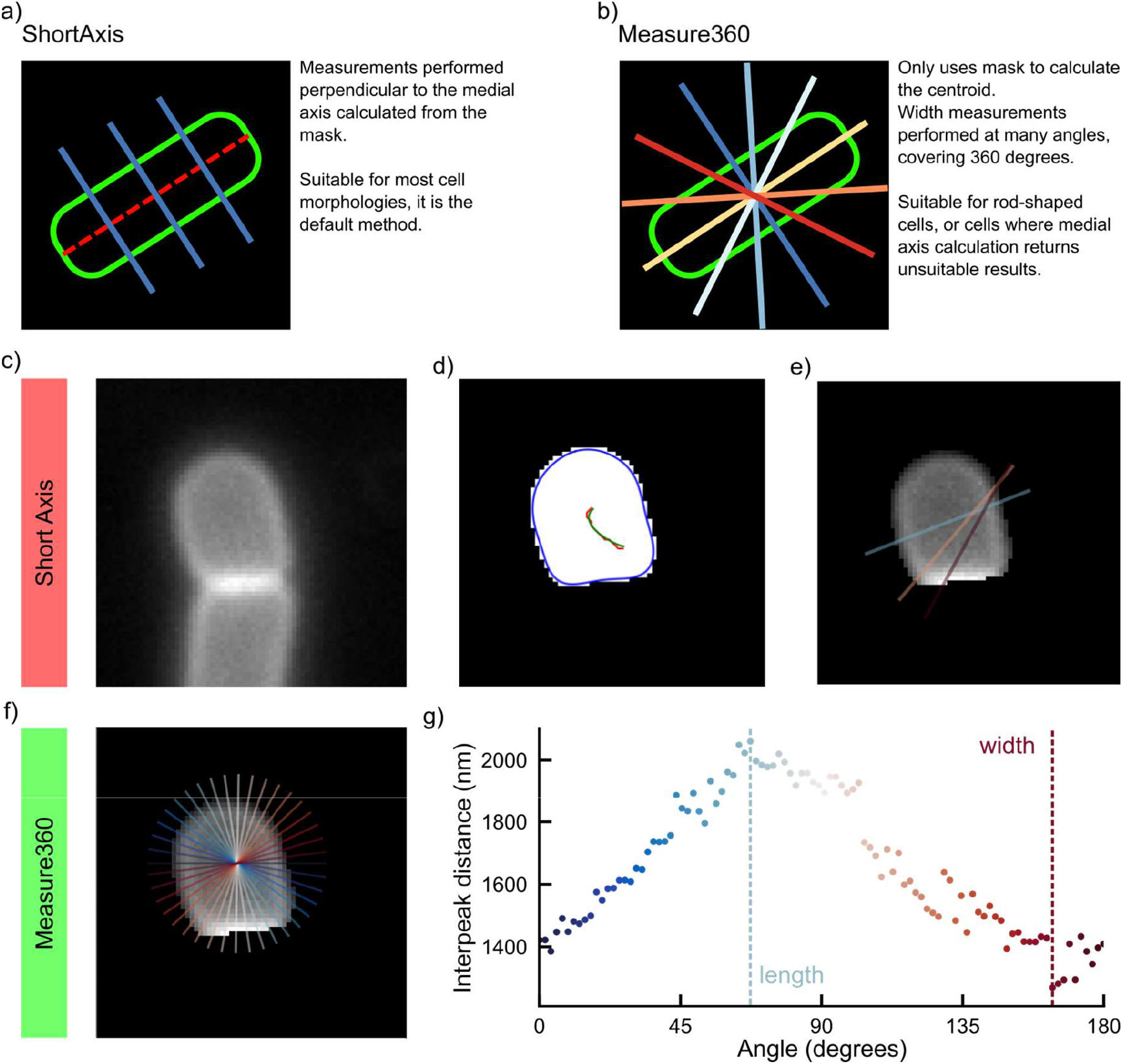
a-b: comparison between the “ShortAxis” and “Measure360” methods for determining lengths and widths of a bacteria cells. c-e: pitfalls of relying on cell midlines to determine intensity profile to fit for width measurements. f-g: Measure360 overcomes the pitfalls of the Short Axis method by fitting intensity profiles at different angles around the cell and then determining width and length as the shortest and longest distances found respectively.

### micromorph enables sensitive detection of width phenotypes

To further validate our width measurement approach, we analysed a dataset from our previously published study on the effects of rodA expression in *Bacillus subtilis* (2). Middlemiss *et al*. found that cells where *rodA* is either over-or under-expressed are wider than cells with native levels of *rodA* expression. This dataset is a good test for our morphology measuring toolkit for a few reasons: (i) the width phenotype is subtle, with differences in median widths being of the order of ∼100nm; (ii) cells exhibit clumping in some cases, thus making traditional segmentation approaches not easily applicable; (iii) cells where *rodA* is under expressed have an unusual shape, closer to a bulging sphere than a rod.

Due to these challenges, in particular due to cell clumping, the images were not analysable with existing automated morphology analysis software. We therefore previously opted for a semi-automated algorithm where line intensity profiles were obtained by manually drawing along the cell short axis and cell width determined by fitting these line profiles with a custom written code.

Here, we reanalysed the dataset using *micromorph* and the Measure360 algorithm. We were not able to analyse the data with the threshold-based segmentation approaches in the other microbial morphology analysis software packages, including *MicrobeJ* and *Oufti*, due to failed segmentation. However, we note that recently (February 2025, v5.13p) *Omnipose* has been integrated into *MicrobeJ*. To make the comparison fair we therefore used the same masks (generated with the *bact_fluo_omni* model *Omnipose* is bundled with) for both software.

Both *micromorph* and *MicrobeJ* with *Omnipose*-generated masks successfully detected the width trend previously observed with semi-manual analysis (Figure 5b). Each analysis method shows a slight offset in the absolute values for cell width compared to the other methods, presumably due to differences in the algorithms and their parameters (Supplementary Figure 2). We investigated this further by comparing the two software’s performance of on our membrane-stained cells synthetic dataset, with results showing *micromorph* was more accurate than *microbeJ* (65.8nm [64.4, 67.0 95% CI] against 86.3nm [84.7, 87.8 95% CI] mean difference). We found *microbeJ* results to be more accurate (mean difference 58.6nm [56.7, 60.4 95% CI]) when masks were generated on low S/N data, which we attribute to the segmentation algorithm producing slightly eroded masks compared to the their high S/N counterparts. On the contrary, *micromorph* measurements were impacted to a lesser degree when low S/N data was used (73.9nm mean difference, [67.7, 80.2 95%CI]). This demonstrates *micromorph* can perform equally to the best-in-class software available on other platforms and can successfully analyse morphology in conditions where software relying on threshold-based segmentation fails.

**Figure 5.**
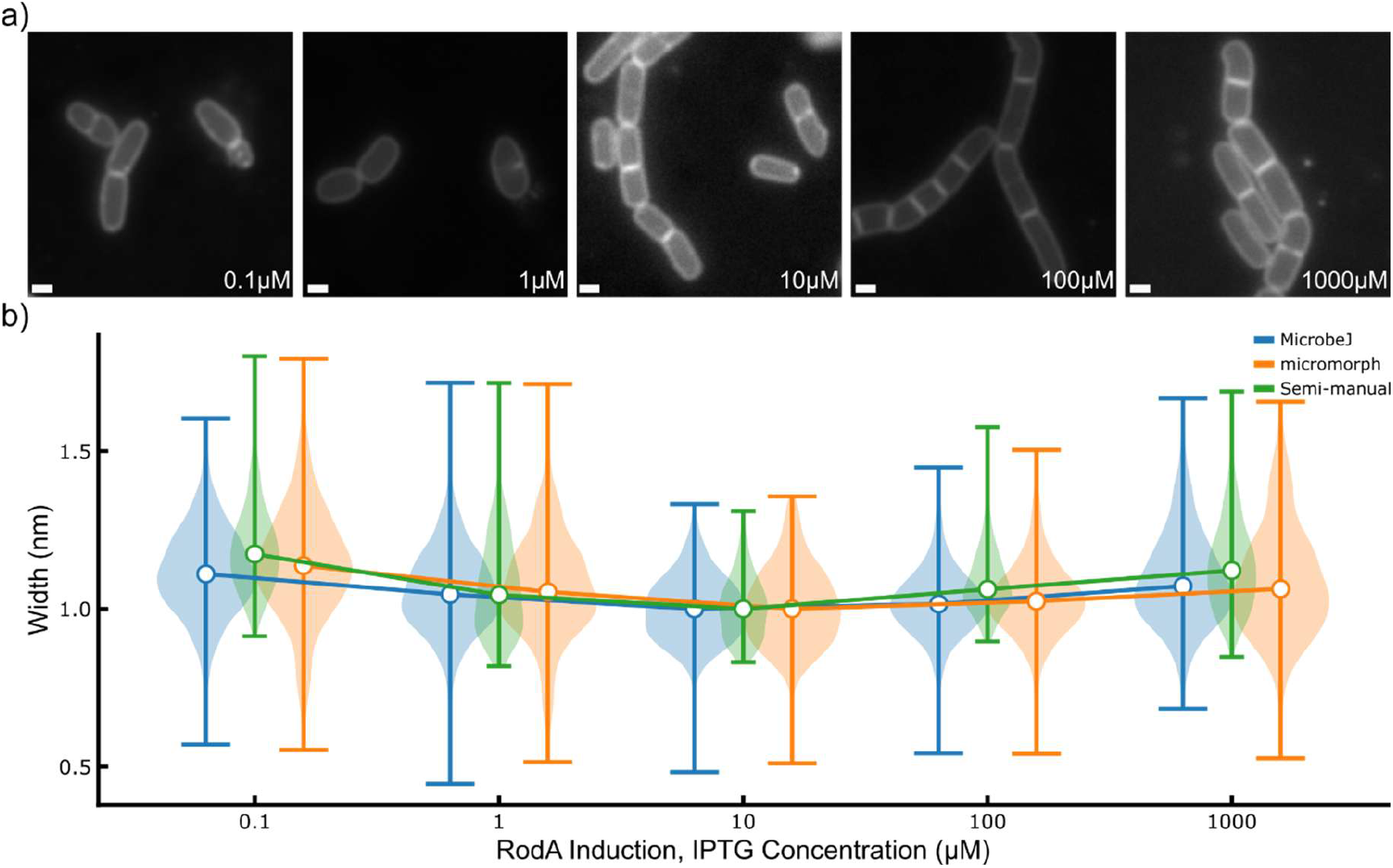
a) Exemplary images for each rodA induction condition of the SM28 (mreB::mreB-HaloTag, Δhag, rodA’:-Pspac-rodA) strain. b) width measurements for all conditions, for each software, normalised with respect to the median width of the 10uM IPTG condition (wild-type rodA induction level). For all cases only widths measured to be within the 800-2000nm range were considered for the analysis. micromorph was able to correctly reproduce the trend found previously by Middlemiss et al., MicrobeJ failed to do so when using threshold-based segmentation methods, but succeeded when using masks generated with Omnipose. Data analysed with Measure360 was filtered with the “Derivative” filter set at 250nm before width finding – see package documentation for more details.

## Testing with different bacterial species demonstrates general applicability of *micromorph*

*micromorph* was developed to be widely applicable and adaptable for custom analysis pipelines. To demonstrate this, we first analysed a dataset of fluorescently labelled *Staph. aureus* cells, which have coccoid shape. We measured the cells widths from phase-contrast and fluorescence microscopy images using the “Measure360” mode. In both cases, we could reliably measure cell widths and lengths, with comparable results between the two measuring modes (Figure 6a-c). The phase-contrast cell width data had a smaller spread than the fluorescence microscopy data (Figure 6b). We attribute this to the fact that the phase contrast microscopy analysis could not differentiate between two dividing cells and a single cell, thus making all such cells appear as one cell. We performed estimation statistics analysis on both width and length datasets, finding that length measurements had 1.66nm mean difference [-8.17, 11.2 95% CI], while width measurements had a mean difference of 102nm [94.7, 110 95% CI] (Figure 6b-c, Supplementary Table 1). We attributed this to the different point spread functions of each imaging modality.

**Figure 6:**
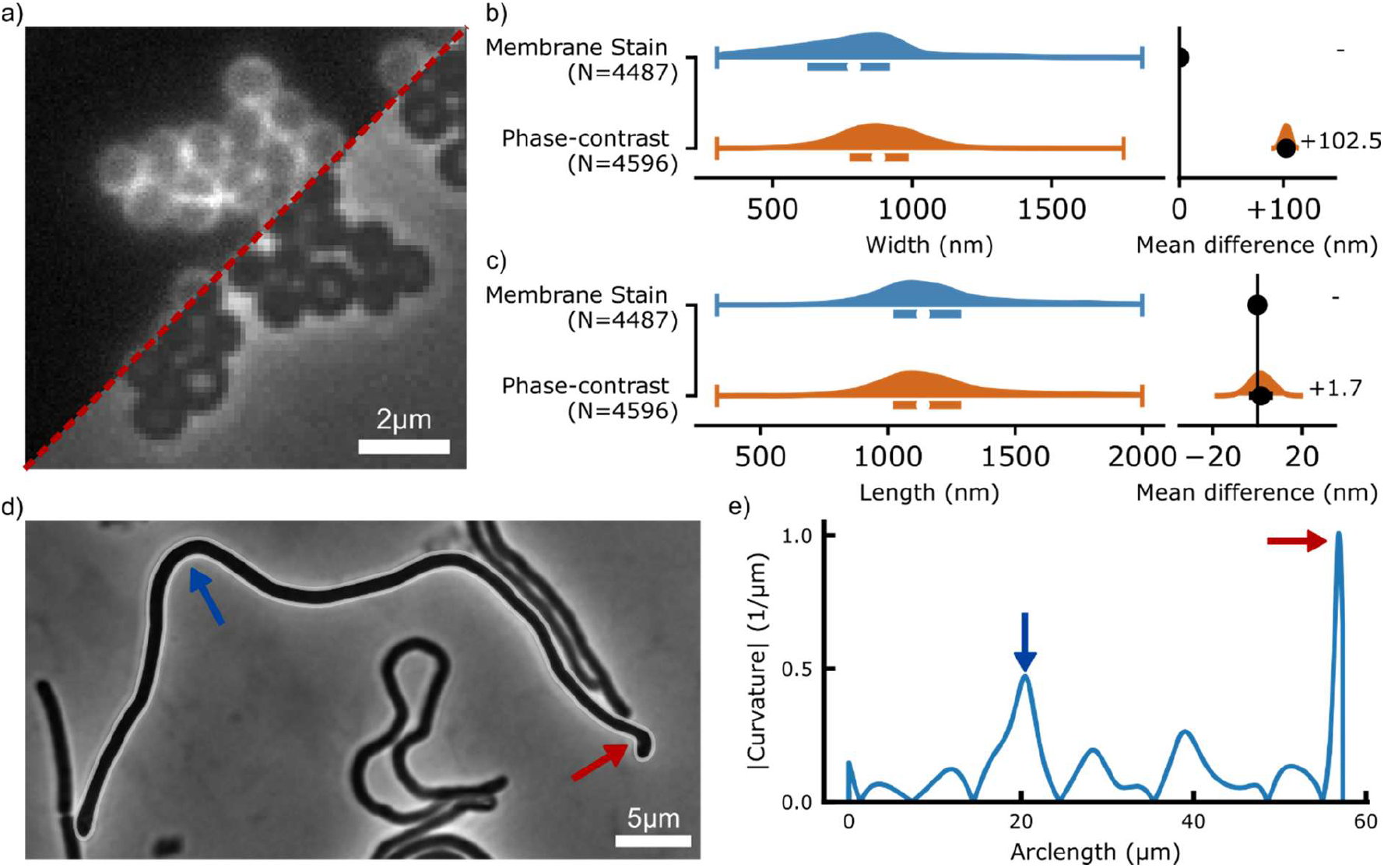
(a) Phase contrast and fluorescence microscopy images of Staph. aureus cells stained with sBADA for 30 minutes. (b-c) Width and length measurements from both image sets, with associated DABEST plots for effect size analysis. (d) Phase contrast image of B. subtilis cells in which ftsW expression has been repressed. White outline is the boundary of the cell used for the curvature analysis showed in (e). Red and blue lines to highlight highest curvature portions of the cell midline.

Second, we analysed data from a *B. subtilis ftsW* CRISPRi knockdown strain. Induction of the CRISPRi system with xylose leads to depletion of *ftsW*, a key cell division protein, leading to cell filamentation. As *micromorph*’s midline and boundary finding algorithm works well for filamentous cells, we could straightforwardly determine the morphological properties of these cells. In order to demonstrate how *micromorph* can easily be extended and integrated into custom data analysis pipelines by taking advantage of its API, we wrote a Jupyter notebook to calculate and identify regions of high curvature in cells (Figure 6d-e). We have made the notebook used for this analysis available in the *micromorph* Github repository.

## Discussion

We have shown *micromorph* is a robust and customisable package for bacterial morphology analysis. *micromorph* works reliably on densely packed cells and for multiple types of microscopy images, namely phase contrast, fluorescence membrane stain and fluorescence cytoplasm label, types of microscopy images. The examples shown here focus on precise measurements of bacterial lengths and widths, but other morphological parameters can also be calculated such as cell area, perimeter and curvature, making *micromorph* suitable for a wide range of bacterial cell biology analyses.

The *micromorph* code is available both as a user-friendly *napari* package and as a well-documented Python package for script based analysis. The *micromorph* Python package will enable advanced users to integrate *micromorph* into their own custom analysis pipelines. For example, functions relating to midline and boundary calculations could be integrated in pipelines aimed at studying the spatiotemporal dynamics of fluorescent proteins of interest. Similarly, users could take advantage of the API to integrate *micromorph* into smart microscopy pipelines for live image analysis and processing.

Areas for future work include functionalities for analysis of time-lapse data and of branched cells. We are also currently investigating integration of *micromorph*’s morphology analysis with single molecule tracking measurements to enable analysis of tracking data within a cellular reference frame, even at high cell density.

## Acknowledgements

We thank Andrew McMahon, David Roberts and Nicholas Briggs for testing early versions of the package. This work was supported by the Wellcome Trust [227452/Z/23/Z to SH].

## Supplementary Information Legend

Supplementary Figure 1: Example of adjacent membrane-stained *B. subtilis* cells.

Supplementary Figure 2: Comparison of *micromorph, microbeJ* and semi-manual cell width measurements

Supplementary Figure 3: Example synthetic data used for benchmarking

Supplementary Figure 4: Benchmark results for membrane-stained synthetic cells

Supplementary Figure 5: Benchmark results for cytoplasmic fluorescent cells

Supplementary Figure 6: Benchmark results for adjacent cells

Supplementary Figure 7: Cell width comparison across different shapes using the ShortAxis and measure360 methods

Supplementary Figure 8: Cell length comparison across different shapes using the ShortAxis and measure360 methods

Supplementary Note 1: Width measurements from intensity profiles

Supplementary Table 1: Effect Sizes

Supplementary Table 2: Summary of Benchmarks

## References

1. Dion MF, Kapoor M, Sun Y, Wilson S, Ryan J, Vigouroux A, et al. Bacillus subtilis cell diameter is determined by the opposing actions of two distinct cell wall synthetic systems. Nat Microbiol. 2019 Aug;4(8):1294–305. doi:10.1038/s41564-019-0439-0

2. Middlemiss S, Blandenet M, Roberts DM, McMahon A, Grimshaw J, Edwards JM, et al. Molecular motor tug-of-war regulates elongasome cell wall synthesis dynamics in Bacillus subtilis. Nat Commun. 2024 Jun 26;15(1):5411. doi:10.1038/s41467-024-49785-x

3. al-Mosleh S, Gopinathan A, Santangelo CD, Huang KC, Rojas ER. Feedback linking cell envelope stiffness, curvature, and synthesis enables robust rod-shaped bacterial growth. Proc Natl Acad Sci. 2022 Oct 11;119(41):e2200728119. doi:10.1073/pnas.2200728119

4. Juillot D, Cornilleau C, Deboosere N, Billaudeau C, Evouna-Mengue P, Lejard V, et al. A High-Content Microscopy Screening Identifies New Genes Involved in Cell Width Control in Bacillus subtilis. Van Wezel GP, editor. mSystems. 2021 Dec 21;6(6):e01017–21. doi:10.1128/mSystems.01017-21

5. Ducret A, Quardokus EM, Brun YV. MicrobeJ, a tool for high throughput bacterial cell detection and quantitative analysis. Nat Microbiol. 2016 Jun 20;1(7):16077. doi:10.1038/nmicrobiol.2016.77 PubMed PMID: 27572972; PubMed Central PMCID: PMC5010025.

6. When Phase Contrast Fails: ChainTracer and NucTracer, Two ImageJ Methods for Semi-Automated Single Cell Analysis Using Membrane or DNA Staining | PLOS One [Internet]. [cited 2025 Apr 13]. Available from: https://journals.plos.org/plosone/article?id=10.1371/journal.pone.0151267

7. van Raaphorst R, Kjos M, Veening JW. BactMAP: An R package for integrating, analyzing and visualizing bacterial microscopy data. Mol Microbiol. 2020;113(1):297–308. doi:10.1111/mmi.14417

8. Jeckel H, Drescher K. Advances and opportunities in image analysis of bacterial cells and communities. FEMS Microbiol Rev. 2021 Jul 1;45(4):fuaa062. doi:10.1093/femsre/fuaa062

9. Archit A, Freckmann L, Nair S, Khalid N, Hilt P, Rajashekar V, et al. Segment Anything for Microscopy. Nat Methods. 2025 Mar;22(3):579–91. doi:10.1038/s41592-024-02580-4

10. Cutler KJ, Stringer C, Lo TW, Rappez L, Stroustrup N, Brook Peterson S, et al. Omnipose: a high-precision morphology-independent solution for bacterial cell segmentation. Nat Methods. 2022 Nov;19(11):1438–48. doi:10.1038/s41592-022-01639-4

11. Panigrahi S, Murat D, Le Gall A, Martineau E, Goldlust K, Fiche JB, et al. Misic, a general deep learning-based method for the high-throughput cell segmentation of complex bacterial communities. Xiao J, Storz G, Hensel Z, editors. eLife. 2021 Sep 9;10:e65151. doi:10.7554/eLife.65151

12. Sen S, Vairagare I, Gosai J, Shrivastava A. RABiTPy: an open-source Python software for rapid, AI-powered bacterial tracking and analysis. BMC Bioinformatics. 2025 May 18;6:127. doi:10.1186/s12859-025-06145-w PubMed PMID: 40383775; PubMed Central PMCID: PMC12087136.

13. Ahmadi A, Dostmohammadi A, Mclean R, Ingalls B. TrackRefiner a tool for refinement of bacillus cell tracking data. Npj Syst Biol Appl. 2025 Nov 17;11(1):127. doi:10.1038/s41540-025-00600-3

14. Sofroniew N, Lambert T, Bokota G, Nunez-Iglesias J, Sobolewski P, Sweet A, et al. napari: a multi-dimensional image viewer for Python [Internet]. Zenodo; 2025 [cited 2025 Apr 15]. Available from: https://zenodo.org/records/15193038 doi:10.5281/zenodo.15193038

15. de Jong IG, Beilharz K, Kuipers OP, Veening JW. Live Cell Imaging of Bacillus subtilis and Streptococcus pneumoniae using Automated Time-lapse Microscopy. J Vis Exp JoVE. 2011 Jul 28;(53):3145. doi:10.3791/3145 PubMed PMID: 21841760; PubMed Central PMCID: PMC3197447.

16. Lambert T. tlambert03/microsim [Python] [Internet]. 2026 [cited 2026 Mar 25]. Available from: https://github.com/tlambert03/microsim

17. Van Der Walt S, Schönberger JL, Nunez-Iglesias J, Boulogne F, Warner JD, Yager N, et al. scikit-image: image processing in Python. PeerJ. 2014 Jun 19;2:e453. doi:10.7717/peerj.453

18. Schindelin J, Arganda-Carreras I, Frise E, Kaynig V, Longair M, Pietzsch T, et al. Fiji: an open-source platform for biological-image analysis. Nat Methods. 2012 Jul;9(7):676–82. doi:10.1038/nmeth.2019

19. Paintdakhi A, Parry B, Campos M, Irnov I, Elf J, Surovtsev I, et al. Oufti: an integrated software package for high-accuracy, high-throughput quantitative microscopy analysis. Mol Microbiol. 2016;99(4):767–77. doi:10.1111/mmi.13264

20. Ho J, Tumkaya T, Aryal S, Choi H, Claridge-Chang A. Moving beyond P values: data analysis with estimation graphics. Nat Methods. 2019 Jul;16(7):565–6. doi:10.1038/s41592-019-0470-3

21. O’Connor OM, Alnahhas RN, Lugagne JB, Dunlop MJ. DeLTA 2.0: A deep learning pipeline for quantifying single-cell spatial and temporal dynamics. PLOS Comput Biol. 2022 Jan 18;18(1):e1009797. doi:10.1371/journal.pcbi.1009797

22. Lo TW, Cutler KJ, Choi HJ, Wiggins PA. OmniSegger: A time-lapse image analysis pipeline for bacterial cells. PLOS Comput Biol. 2025 May 28;21(5):e1013088. doi:10.1371/journal.pcbi.1013088

23. Blum H. A Transformation for Extracting New Descriptors of Shape. MIT Press. 1967;362–80.

24. Whitley KD, Jukes C, Tregidgo N, Karinou E, Almada P, Cesbron Y, et al. FtsZ treadmilling is essential for Z-ring condensation and septal constriction initiation in Bacillus subtilis cell division. Nat Commun. 2021 Apr 27;12(1):2448. doi:10.1038/s41467-021-22526-0

