## Supplementary Information for "*micromorph*: a Python toolkit for measurement of microbial morphology"

### Supplementary Materials

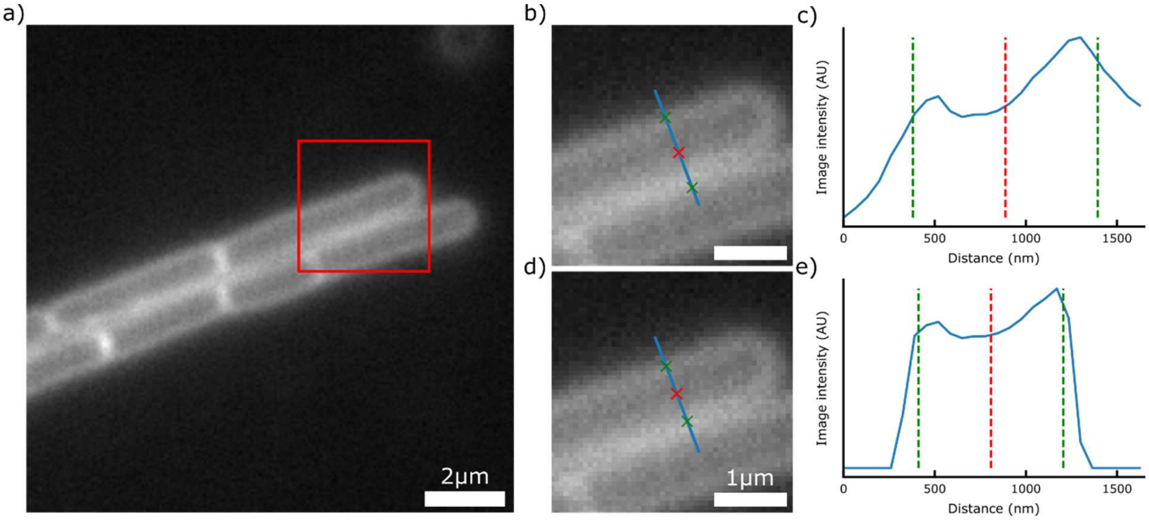


*Supplementary Figure 1 a) example image of membrane-stained B. subtilis cells. Red box indicates area zoomed in panels (b, d), where the blue line indicates where the profile was taken from and the red and green crosses indicate the cell centre and edges respectively. (c, e) Line profiles with fit results for the case in which the original image is used (c) and the case where the mask is applied to the image before measuring the intensity profile (e). Without mask application, the cell centre is incorrectly placed too close to the adjacent cell, and a higher width value is measured (887nm against 808nm when the mask is applied).*

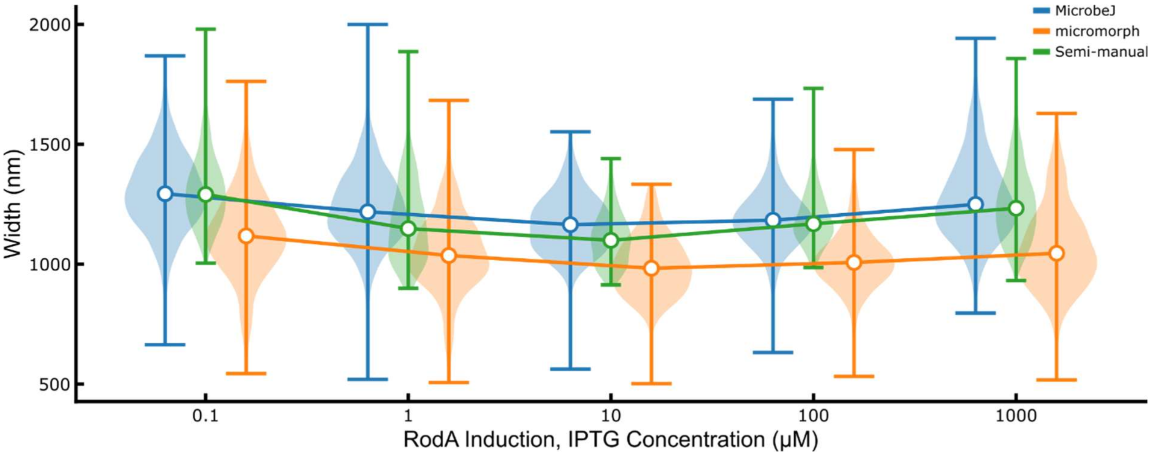


*Supplementary Figure 2 Width measurements for the strain described in Figure 5. micromorph was able to successfully detect the width trend, while MicrobeJ only succeeded when segmenting cells using Omnipose. Offsets in absolute values measured from different software are expected due to the specifics of each algorithm.*

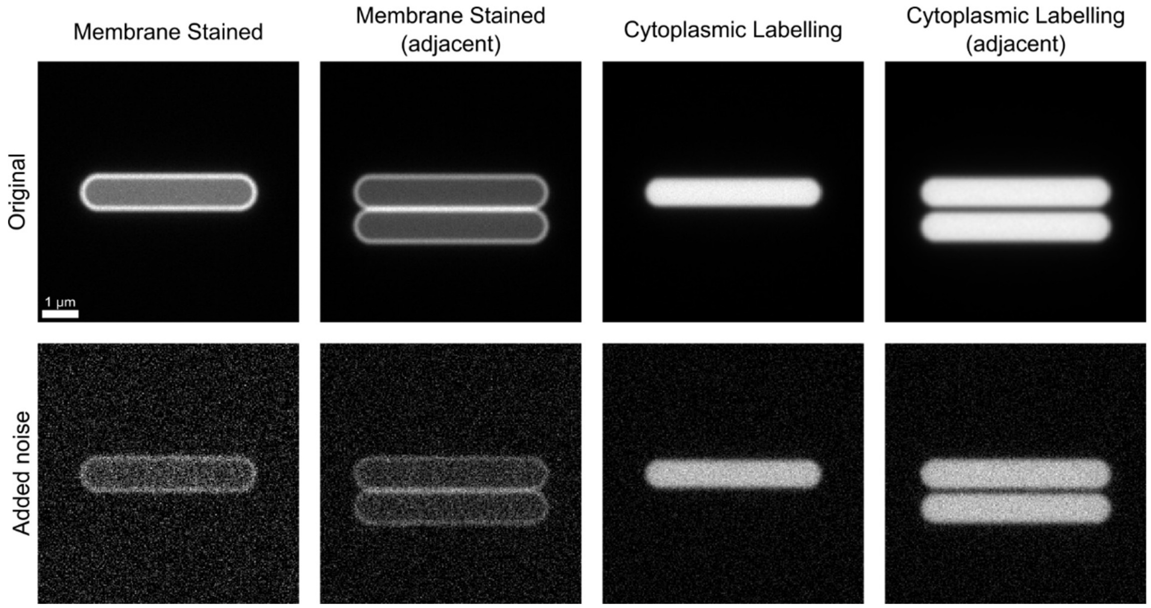


*Supplementary Figure 3 Example synthetic images generated for benchmarking the width finding algorithms for membrane stained and cytoplasmic labelled/phase-contrast cells – see Supplementary Table 2 for benchmark results and Methods for details on image generation*

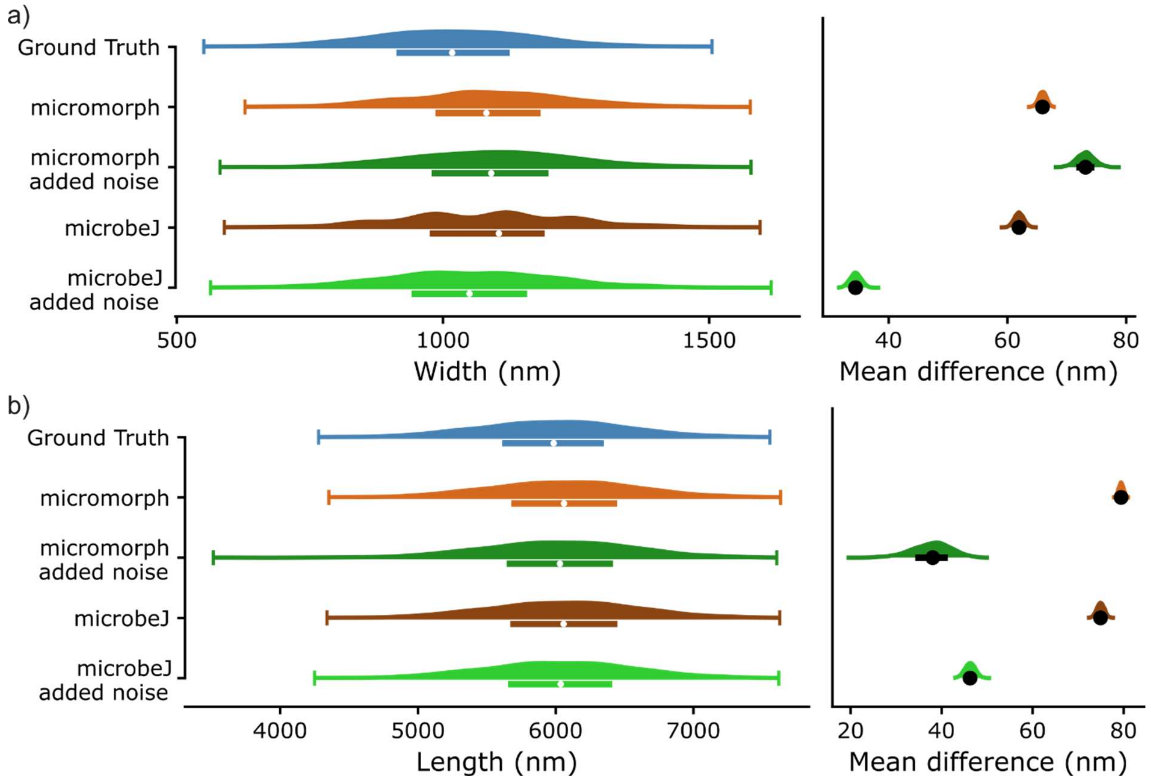


*Supplementary Figure 4 Benchmark results for the membrane-stained synthetic data, using* micromorph and microbeJ.

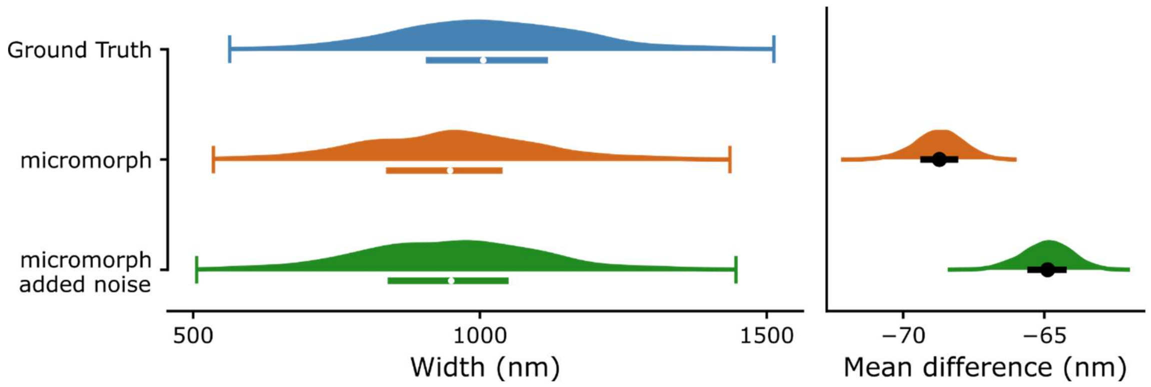


*Supplementary Figure 5 Benchmark results for the width finding algorithm for cells with cytoplasmic fluorescence.*

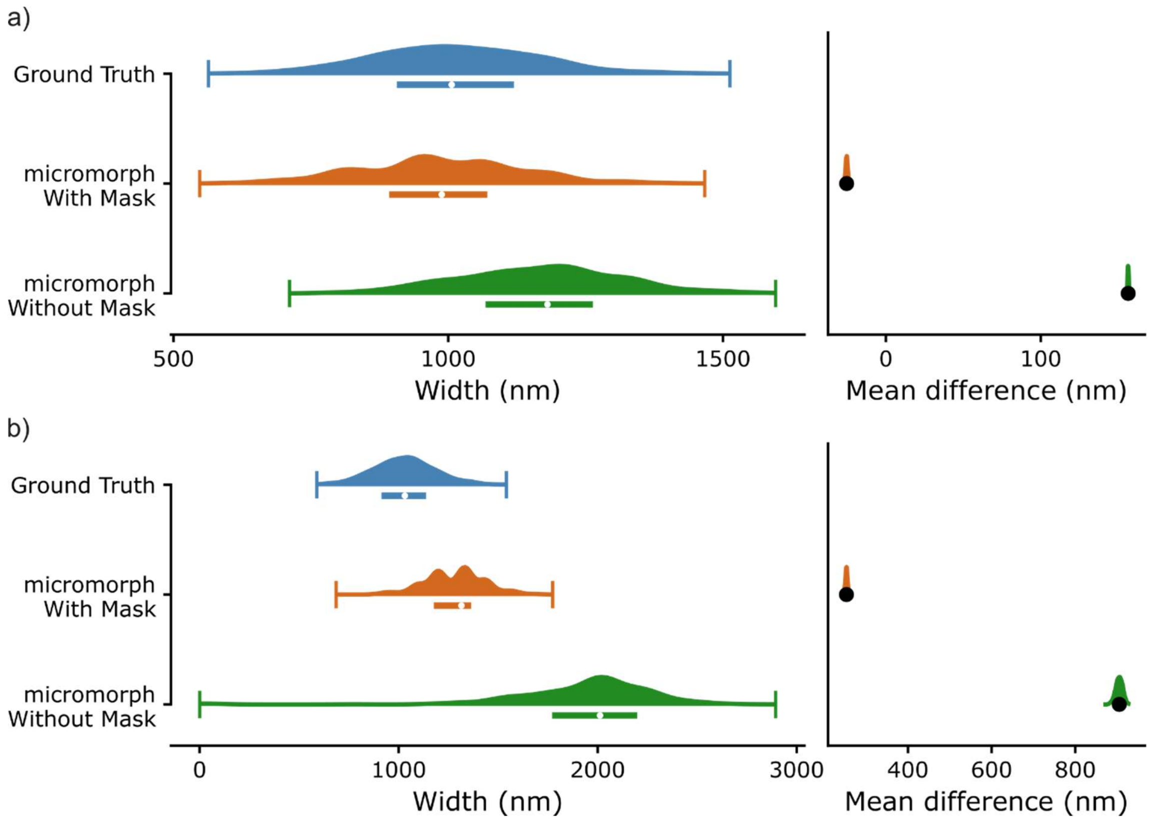


*Supplementary Figure 6 Benchmark results for cells adjacent to each other, with and without mask application for the case of membrane-stained cells (a) or cytoplasmic labelled cells (b)*

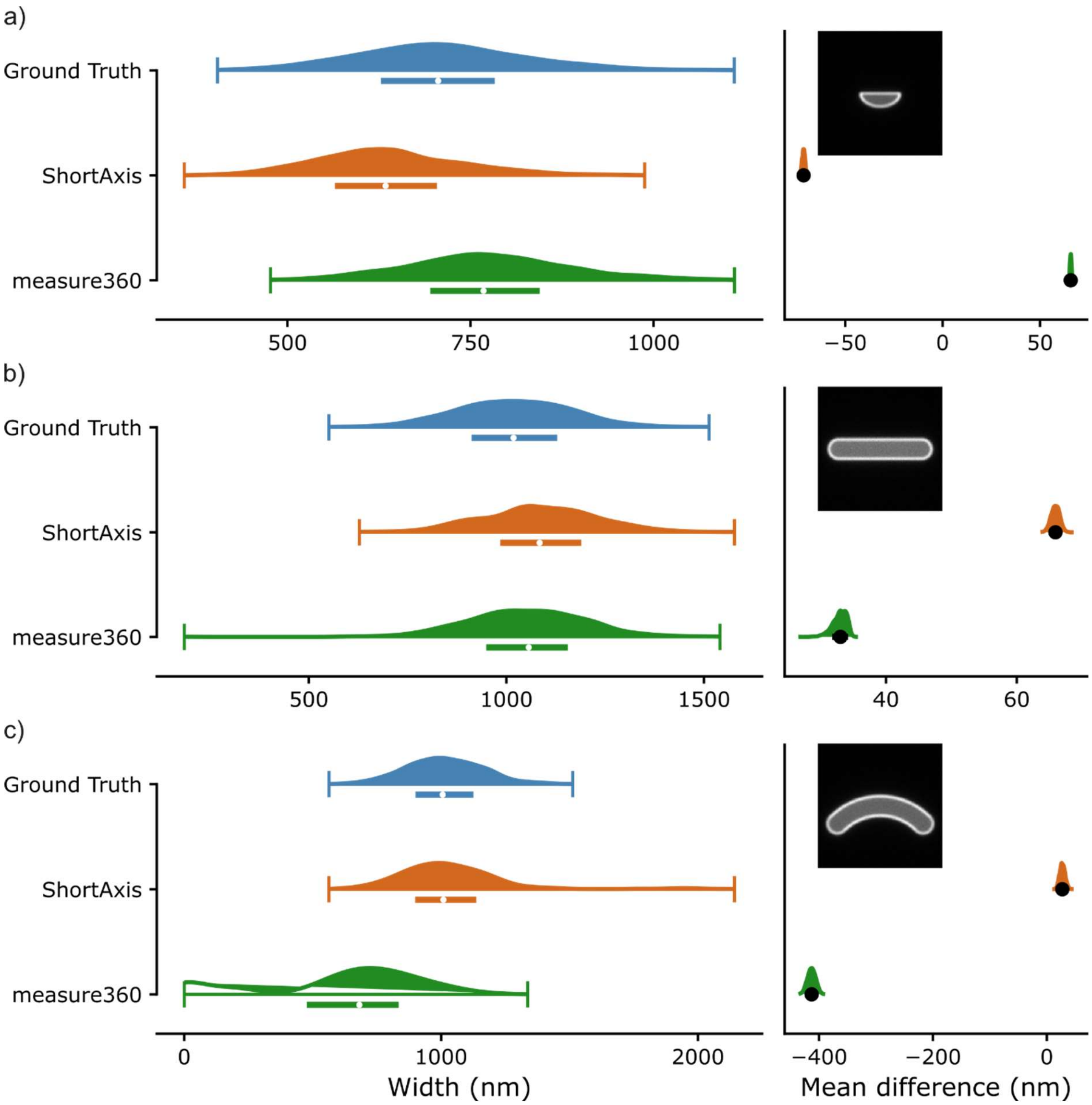


*Supplementary Figure 7 Cell width analysis for three types of cell shapes – half moon (a), rod-shaped (b) and elongated (c) – using both the ShortAxis and measure360 methods.*

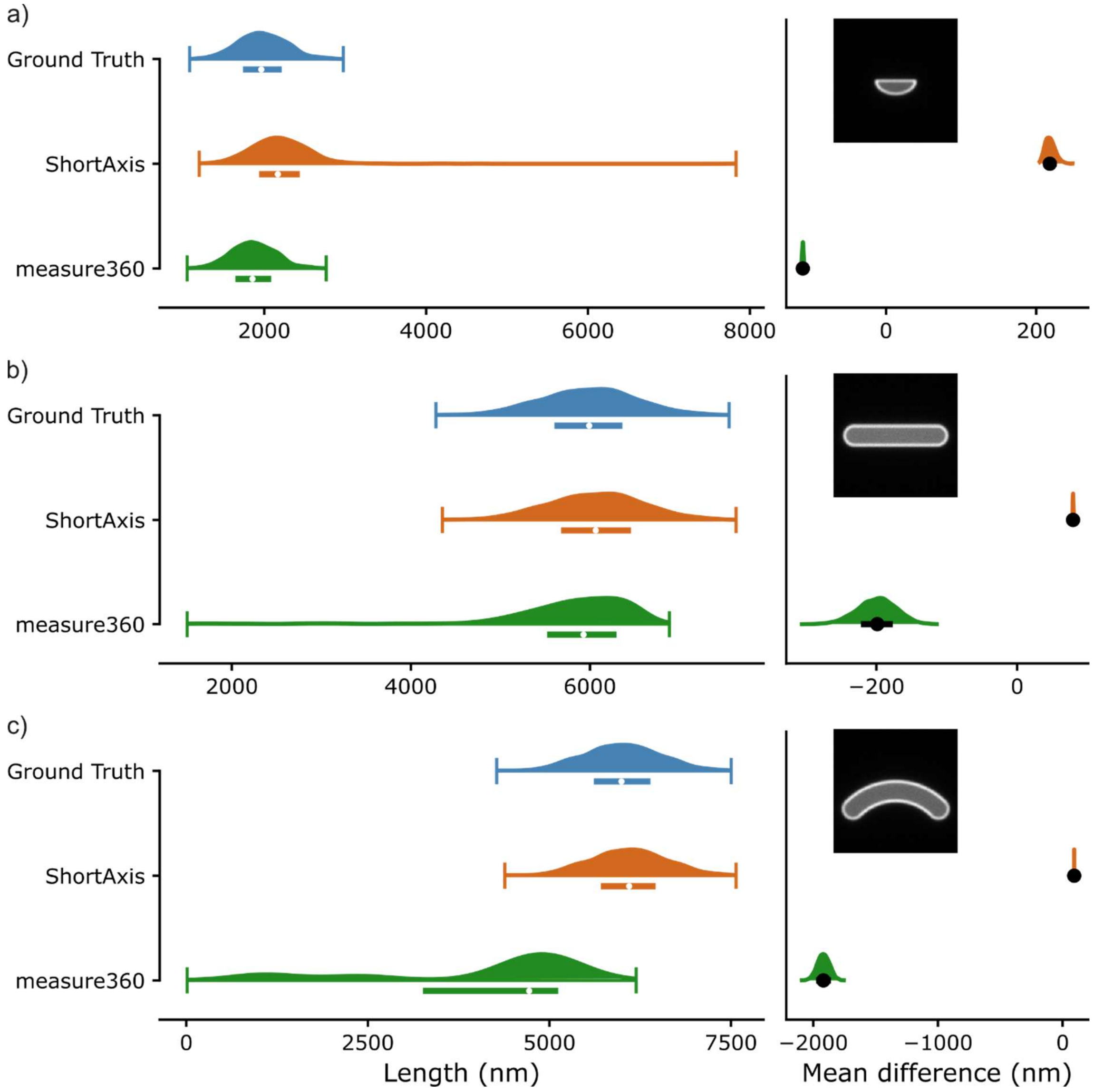


*Supplementary Figure 8 Cell length analysis for three types of cell shapes – half moon (a), rod-shaped (b) and elongated (c) – using both the ShortAxis and measure360 methods.*

#### *Width measurements from intensity profiles*

Three fit options are available to extract widths, or more generally, distances, from intensity profiles.

The first method is based on the tilted-ring profile model described in Whitley et al (24). Consider a fluorescently labelled circle of radius $r$. Our aim is to model the intensity profile of the circle projected on the x-axis.

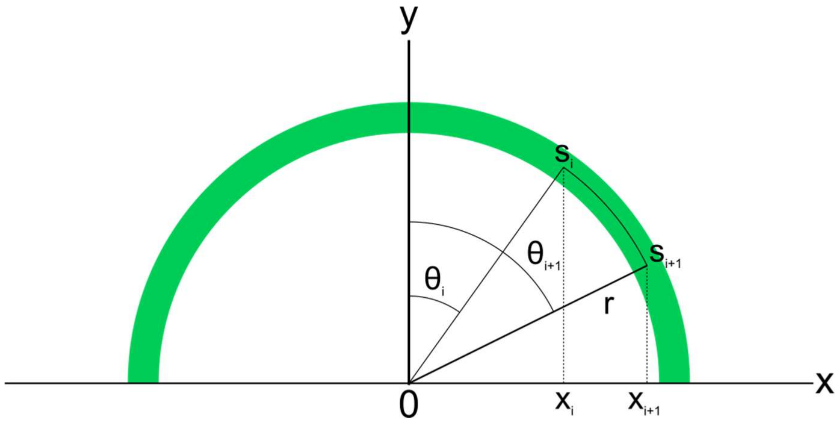


If we consider only half of the circle, the fluorescence intensity between two points xi and xi+1 will be proportional to the arclength segment directly above it:

$$I(x_i) = I_0(s_{i+1} - s_i)$$

Where $I_0$ is the intensity per unit length of the circle. Since $x_i = r\sin\theta_i$ we can express $s_i$ as

$$s_i = r\sin^{-1}(\frac{x_i}{r})$$

So that the arclength difference is

$$s_{i+1} - s_i = r(\sin^{-1}\left(\frac{x_{i+1}}{r}\right) - \sin^{-1}\left(\frac{x_i}{r}\right))$$

Which means the full intensity measured in segment $i$ is

$I(x_i) = 2I_0 r(\sin^{-1}\left(\frac{x_{i+1}}{r}\right) - \sin^{-1}\left(\frac{x_i}{r}\right))$The second method is used to fit phase contrast images. In this case, an inverted top-hat profile (or super-gaussian) is fitted to the intensity profile:

$$I(x) = I_0 + A * \exp\left(-2 * \left({}^{x - x_0}/_{w}\right)^{\alpha}\right)$$

Where $I_0$ is the background intensity, $A$ the amplitude, $x_0$ the centre of the gaussian, $w$ its FWHM (full width half maximum) and $\alpha$ an exponent which controls the top-hat nature of the profile. We have found it best to keep $\alpha$ fixed to a value of 4 for good fitting, but this could be changed by users.

In the third case, we use the same distribution (but inverted) to fit cytoplasmic signal.

**Supplementary Table 1: Effect Sizes**

| Figure number | Quantity, comparison | Difference | 95% CI | No. of data points (N1, N2) |
|---|---|---|---|---|
| 2g | *Cell width,* membrane-stained minus cytoplasmic GFP | 28.9nm | [17.9, 39.4] | (1227, 1242) |
| | *Cell width,* membrane-stained minus phase-contrast | 24.9nm | [13.4, 36.9] | (1227, 1234) |
| 6b | *Cell width, "*membrane-stained" minus "phase-contrast" | 102nm | [94.7, 1.1e+02] | (4487, 4596) |
| | Cell length, "membrane-stained" minus "phase-contrast" | 1.66nm | [-8.17, 11.2] | (4487, 4596) |

**Supplementary Table 2: Summary of Benchmarks**

| Supplementary Figure number | Test Dataset | Quantity, comparison | Difference | 95% CI | No. of data points (N1, N2) | Mean Absolute Error |
|---|---|---|---|---|---|---|
| 4a | Rod-shaped, membrane-stained | *Cell width* – ground truth minus measurement, no noise | 65.8nm | [64.4, 67.0] | (1000, 1000) | 67.41nm |
| 4a | | *Cell width* – ground truth minus measurement, added noise | 73.9nm | [67.7, 80.2] | (1000, 1000) | 85.86nm |
| 4b | | *Cell length* – ground truth minus measurement, no noise | 90.2nm | [88.8, 91.6] | (1000, 1000) | 90.2nm |
| 4b | | *Cell length* – ground truth minus measurement, added noise | 37.8nm | [26.3, 44.4] | (1000, 1000) | 64.9nm |
| 4b | | *Cell width* – ground truth minus measurement, no noise (microbeJ) | 86.3nm | [84.7, 87.8] | (1000, 1000) | 86.2nm |
| 4a | | *Cell width* – ground truth minus | 58.6nm | [56.7, 60.4] | (1000, 1000) | 59.1nm |

| | | | | | | |
|---|---|---|---|---|---|---|
| | | measurement, added noise (microbeJ) | | | | |
| 4b | | *Cell length* – ground truth minus measurement, no noise (microbeJ) | 69.1nm | [64.6, 71.6] | (1000, 1000) | 72.8nm |
| | | *Cell length* – ground truth minus measurement, added noise (microbeJ) | 46.2nm | [43.8, 48.5] | (1000, 1000) | 51.0nm |
| 5 | Rod-shaped, cytoplasmic labelling. | *Cell width* – ground truth minus measurement, no added noise | -67.8nm | [-69.8, -65.2] | (1000, 1000) | 69.2nm |
| 5 | | *Cell width* – ground truth minus measurement, added noise | -64.9nm | [-66.9, -63.3] | (1000, 1000) | 64.8nm |
| 6a | Rod-shaped, membrane-stained. Adjacent cells. | *Cell width* – ground truth minus measurement, mask off | -25.1nm | [-26.5, -23.7] | (1000, 1000) | 28.2nm |
| 6a | | *Cell width* – ground truth minus measurement, mask on | 156nm | [156, 157] | (1000, 1000) | 156nm |
| 6b | Rod-shaped, cytoplasmic labelling. Adjacent cells. | *Cell width* – ground truth minus measurement, mask off | -138nm | [-140, -137] | (1000, 1000) | 139nm |
| 6b | | *Cell width* – ground truth minus measurement, mask on | 870nm | [844, 890] | (1000, 1000) | 908 |

| | | | | | | |
|---|---|---|---|---|---|---|
| 7a | Half-moon, membrane-stained. | *Cell width* – ground truth minus measurement (measure360, half-ellipse) | 65.8nm | [65.0, 66.5] | (1000, 1000) | 65.9 |
| 7a | | *Cell width* – ground truth minus measurement (short axis, half-ellipse) | -71.4nm | [-72.9, -70.0] | (1000, 1000) | 71.6 |
| 8a | | *Cell length* – ground truth minus measurement (measure360, half-ellipse) | -1.11e+02nm | [-1.13e+02, -1.09e+02] | (1000, 1000) | 110.9 |
| 8a | | *Cell length* – ground truth minus measurement (short axis, half-ellipse) | 2.19e+02nm | [2.09e+02, 2.41e+02] | (1000, 1000) | 218.6 |
| 7b | Rod-shaped, membrane-stained. | *Cell width* – ground truth minus measurement (measure360, rod-shaped) | 71.1nm | [69.5, 72.7] | (1000, 1000) | 35.2 |
| 7b | | *Cell width* – ground truth minus measurement (short axis, rod-shaped) | 65.8nm | [64.4, 67.0] | (1000, 1000) | 67.41nm |
| 8a | | *Cell length* – ground truth minus measurement (measure360, rod-shaped) | -1.11e+02nm | [-1.13e+02, -1.09e+02] | (1000, 1000) | 211.9 |
| 8b | | *Cell length* – ground truth minus measurement (short axis, rod-shaped) | 90.2nm | [88.9, 91.6] | (1000, 1000) | 90.1 |

| | | | | | | |
|---|---|---|---|---|---|---|
| 7c | Elongated, membrane-stained. | *Cell width* – ground truth minus measurement (measure360, elongated) | -4.13e+02 | [-4.28e+02, -3.99e+02] | (1000, 1000) | 412.8 |
| 8c | | *Cell width* – ground truth minus measurement (short axis, elongated) | 27.5nm | [19.2, 37.9] | (1000, 1000) | 38.6 |
| 7c | | *Cell length* – ground truth minus measurement (measure360, elongated) | -1.92e+03nm | [-2.01e+03, -1.82e+03] | (1000, 1000) | 1916.99 |
| 8c | | *Cell length* – ground truth minus measurement (short axis, elongated) | 92.6nm | [91.8, 93.5] | (1000, 1000) | 92.6 |
